# Approaching pathogenic *Clostridia* from a One Health perspective

**DOI:** 10.1101/2024.01.08.574718

**Authors:** Laura M. Cersosimo, Jay N. Worley, Lynn Bry

## Abstract

Spore-forming pathogens have a unique capacity to thrive in diverse environments, and with temporal persistence afforded through their ability to sporulate. These behaviors require a One Health approach to identify critical reservoirs and outbreak-associated transmission chains, given their capacity to freely move across soils, waterways, foodstuffs, and as commensals or infecting pathogens in human and veterinary populations. Among anaerobic spore-formers, genomic resources for pathogens including *C*. *botulinum*, *C*. *difficile*, and *C*. *perfringens* enable our capacity to identify common and unique factors that support their persistence in diverse reservoirs and capacity to cause disease. Publicly available genomic resources for spore-forming pathogens at NCBI’s Pathogen Detection program aid outbreak investigations and longitudinal monitoring in national and international programs in public health and food safety, as well as for local healthcare systems. These tools also enable research to derive new knowledge regarding disease pathogenesis, and to inform strategies in disease prevention and treatment. As global community resources, the continued sharing of strain genomic data and phenotypes further enhances international resources and means to develop impactful applications. We present examples showing use of these resources in surveillance, including capacity to assess linkages among clinical, environmental, and foodborne reservoirs and to further research investigations into factors promoting their persistence and virulence in different settings.

## Introduction

The anaerobic *Clostridia* include multiple toxigenic, disease-causing species, particularly *Clostridium botulinum, Clostridioides difficile* and *Clostridium perfringens*^1,2^, in addition to other species such as *Clostridium tetani, Clostridium sordellii* and *Clostridium novyi*^3,4^. A core feature among the *Clostridia* is their ability to sporulate, a stress-induced response that enables their survival through harsh conditions including extremes of temperature, pH, as well as exposures to antimicrobials and other sterilants^5-7^. *Clostridial* endospores are found ubiquitously in marine and freshwater ecosystems, soil, and food sources, in addition to their colonization of the gastrointestinal tracts of animals^8-10^. *Clostridial* spores can further persist for extended periods of time and be actively spread through adherence to fomites such as walls, floors, soles of shoes, animal fur, as well as on surfaces and equipment^11,12^. In contrast to pathogenic strains from other phyla, toxigenic and spore-forming pathogens have greater capacity to spread and persist in diverse environments^12-15^, factors that provide unique challenges in detecting transmission chains, and in considering preventive and therapeutic approaches for infections, particularly in immunocompromised and other vulnerable populations such as infants and young animals^16-19^.

Globally, *Clostridial* pathogens incur billions of US dollars annually in costs associated with human and veterinary infections, as well as agricultural systems and components of the food supply that are impacted by associated outbreaks^20-22^. The advent of methods to rapidly sequence bacterial genomes has provided a robust tool to enhance our monitoring and implementation of preventive programs^23-25^, while also supporting mechanistic studies to define how toxigenic strains leverage these virulence factors to promote their survival and spread among ecosystems^26-29^. Investigating transmission chains among these settings under a One Health framework supports ongoing efforts in Public Health to define pathogen reservoirs, at-risk populations, and the development of preventive programs and improved measures to diagnose, treat, and prevent these infections.

### One Health genomic surveillance for global pathogens

One Health approaches aim to collaboratively improve the health of human and animal populations, and the associated environments that they inhabit. In the case of infectious diseases, it promotes a collaborative approach among diverse disciplines to investigate pathogens that occur across species and ecosystems^30^.

Pathogen genomic analyses have bolstered One Health surveillance efforts for a wide range of diseases^31^, including foodborne illnesses^32^, hospital-acquired infections^33-35^, and zoonoses^36^. Resources for the deposition and monitoring of antibiotic resistance and potential outbreaks are critical components of global genomic surveillance^37,38^. Strain genomic information, alone, cannot define outbreaks or transmission chains, and is most effectively leveraged within surveillance and outbreak detection programs that incorporate robust epidemiologic and statistical analyses to confirm associations among case clusters with outbreaks, and to identify reservoirs from environmental, foodborne, or clinical settings^34,39,40, 23^. As such, means to make strain genomic data publicly available aids ongoing and future efforts to detect pathogen strains as they occur.

NCBI’s Pathogen Detection resource (PD; https://www.ncbi.nlm.nih.gov/pathogens/) supports global genomic surveillance of pathogens from diverse origins, including from foodstuffs, environmental sources, and interactions with host species, including cases of symptomatic infections and of asymptomatic carriage. PD’s Pathogen Genomic Analysis Pipeline (PGAP)^32,41,42^ annotates user-communicated genomic information, returning annotated genes and proteins. The Pathogen Detection pipeline further performs whole genome Multi-Locus Sequence Typing (wgMLST) on communicated genomic data for clustering, with calculation of single nucleotide polymorphism (SNP) distances among isolates, to place strains within global clusters. This placement aids to assess strain relatedness and, relative to the breadth and depth of globally communicated genomes, can place strains within a global context^31,43^. PD resources for *C. difficile* and *C. perfringens,* also currently include well-described toxins for these strains and a framework to incorporate toxin gene information for other toxigenic *Clostridia*. With the AMRFinderPlus tools^42,44^, the PD resources assesses carriage of antibiotic resistance genes, point mutations, stress response, and virulence genes.

User-provided metadata with isolate genomes substantially enhances use of the genomic surveillance efforts and capacity to evaluate strain origin and transmission chains across reservoirs. In particular, datapoints such as "isolate type" capture whether isolates were obtained from clinical, environmental, or other settings. Sub-specifying information, such as site of origin from foodstuffs, soil, or waterway locations, and sub- of clinical-origin strains by anatomic site and whether causative of disease or associated with asymptomatic colonization also enhances the context in which closely related strains have occurred. The One Health Enteric (OHE) package provides open source tools to aid in strain genomic and metadata collection^45^.

Isolate location data, which ranges from information provided at the level of country of origin to municipalities and geo-coordinates, also enhances strain tracking and capacity to provide context for newly contributed genomes. Incorporation of codified clinical data, such as ICD.10 (International Classification of Diseases 10th revision), Diagnosis Related Group (DRG) codes^46,47^, and ordered testing that identified isolates, particularly to distinguish surveillance testing for asymptomatic carriage of toxigenic strains versus association with active infections^48^, provides critical information when tracking strains over time as well as within and among sites and healthcare institutions. Under HIPAA (Health Insurance Portability and Accountability Act)^49^, healthcare institutions in the US are limited from providing full location and date/timestamps that are associated with clinical events, but with institutional review and approval may communicate anonymous or limited datasets (LDS) that note the country, state, and often city, first 2-3 digits of the zip code, relative to population density, and the strain year of origin^33,49,50^.

Additional strain phenotypic data, including antibiograms, biochemical, or serologic strain typing also enhances available datasets to provide functional assessments of antibiotic resistance and known antigenic or biochemical properties^23,47,51^. The phenotypic analyses, particularly when conducted under a recognized international standards such as CLIA (Clinical Laboratory Improvement Amendments)^52^ or ISO 15189^53^, offer means to correlate phenotypic findings from strains collected across geographic locations with genomic findings^39^.

A number of global programs leverage the PD resources and repository^32^, including the US FDA’s GenomeTrakr program^23,35,54,55^, CDC’s PulseNet^37,56^, country-wide and state/province Public Health programs^23,57-59^, and surveillance programs within hospital networks^25,48,56,60,61^. Additional international consortia that incorporate surveillance for toxigenic *Clostridia* include the International Pathogen Surveillance Network (IPSN)^62^, formed in 2023 by the World Health Organization Hub for Pandemic and Epidemic Intelligence to advance use of pathogen genomics in informing decisions made in public health, while improve upstream events in sample collection and for downstream data analysis, and the Wellcome Trust-founded Surveillance and Epidemiology of Drug-resistant Infections Consortium (SEDRIC)^63^.

### C. botulinum, C. difficile and C. perfringens’ genomic resources

Among PD’s pathogen genomic resources, those for the *Clostridia* represent newer additions with the first having been for *C. difficile^48^,* followed by *C. perfringens^23^* and *C. botulinum^64^* (Figure 1, Supplemental Data Files 1 and 2). Figure 1 shows the global distribution of strains contributed as of December 2023, with a majority of genome-sequenced isolates originating from North America, Europe, China, and Australia, and nominal coverage of strain diversity that originates from Africa, the Middle East, Central Asia, insular regions of southeast Asia, and regions within Central and South America. Such maps illustrate where strain coverage may provide denser sampling of underlying genomic diversity to support more in-depth outbreak investigations, while also highlighting global needs and opportunities to enable genomic surveillance efforts in sparsely covered regions.

**Figure 1:**
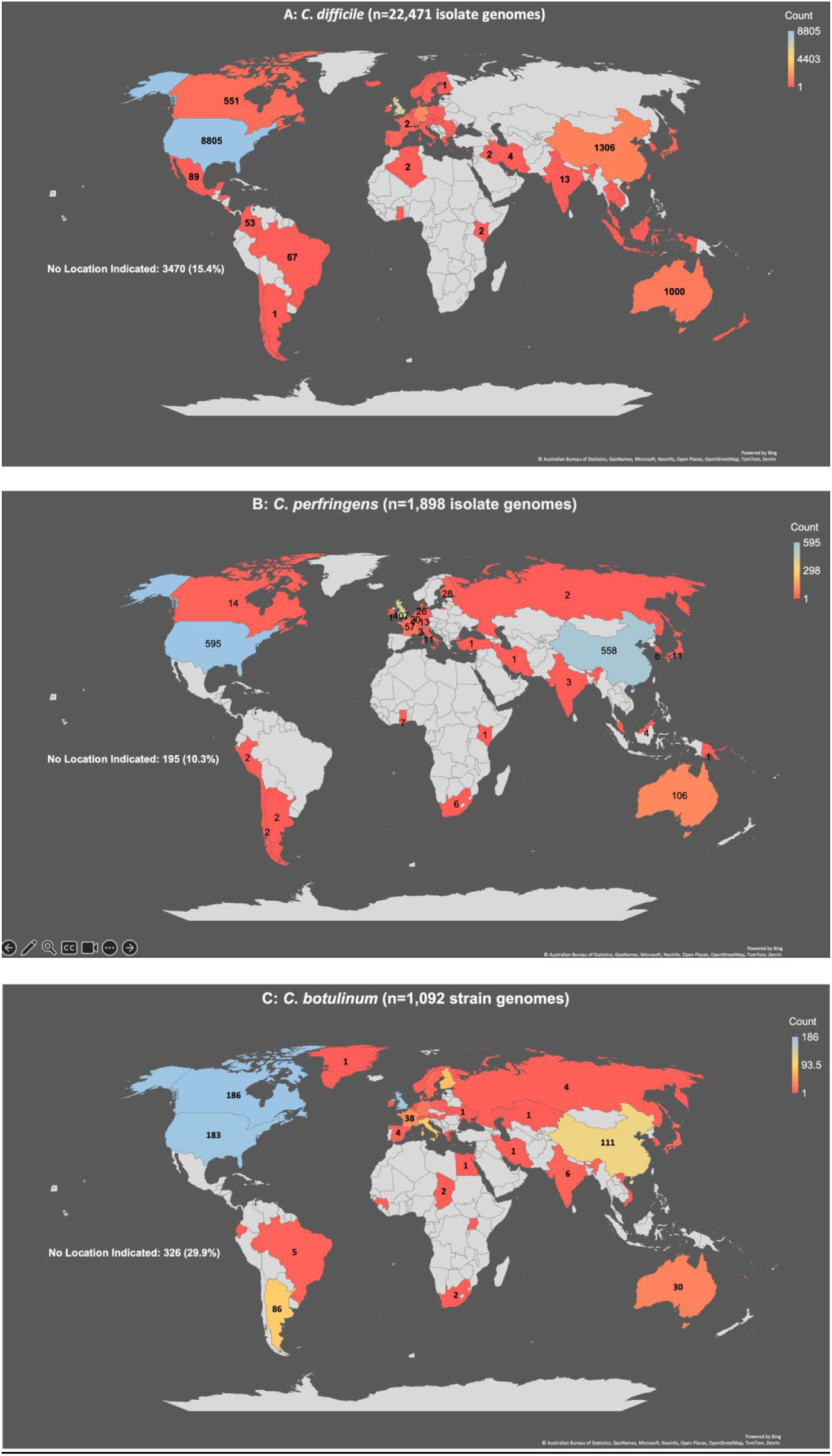
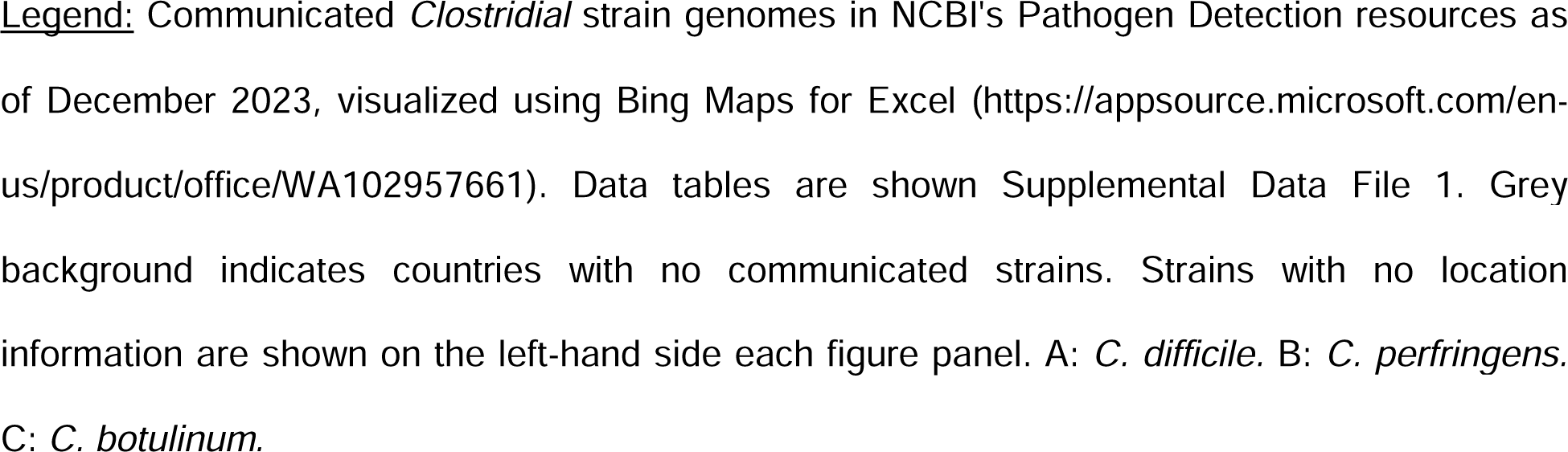
Globally communicated cases of toxigenic *C. difficile, C. perfringens,* and *C. botulinum*.

The distribution of *C*. *difficile* by depositor-reported isolation type (Fig. 1A) is 81% clinical, 2% environmental, and 17% with no isolation type provided. Within the *C*. *perfringens* (Fig. 1B) and *C*. *botulinum* resources (Fig 1C), 56% and 31% of isolates, respectively, originated from clinical sources, and 17% and 53%, respectively from non-clinical samples. Among the 284 environmental isolates of *C. botulinum,* 37% of isolates originated from soil and 35% from food sources. Of the *C*. *perfringens* isolates categorized as environmental samples (n=354), 81% of isolates originated from food sources (26%), fecal (27%), and soil (28%) samples. In contrast, one quarter of environmental *C*. *difficile* isolates originated from soil and 11% from animal fecal samples, with the largest being associated with human clinical isolates in infected patients or asymptomatic carriers. Twenty-five percent of *C*. *difficile* environmental isolates were also characterized as “environmental samples” under isolation source with one geographical location listed demonstrating the need for more descriptive metadata.

### *Clostridial* genomic analysis of virulence factors and antibiotic resistance genes

*Clostridial* strain genomic analyses have been essential to establish the range of toxin systems found within pathogen species, including the large Clostridial toxins and additional factors that contribute to host disease. This latter category includes bacterial genes that may support aspects of innate *Clostridial* metabolism, via excreted proteases, lipases, and other hemolysins that acquire nutrients from the extracellular milieu. However, when expressed in sterile body sites, these factors can promote the development of gas gangrene and other toxemic manifestations of disease^1,65^.

Pathogen Detection analyses of antibiotic resistance gene repertoires have been employed to aid in surveillance and identify risks for triggering disease, particularly for *C. difficile* colitis. One hundred and seventy-one AMR genes and gene-level mutations have been identified in the PD Clostridial resources representing associated resistance to eighteen different classes of antimicrobials (Table 1). Of interest, particular forms of resistance gene carriage suggest interesting insights regarding antibiotic exposures for each species.

**Table 1:**
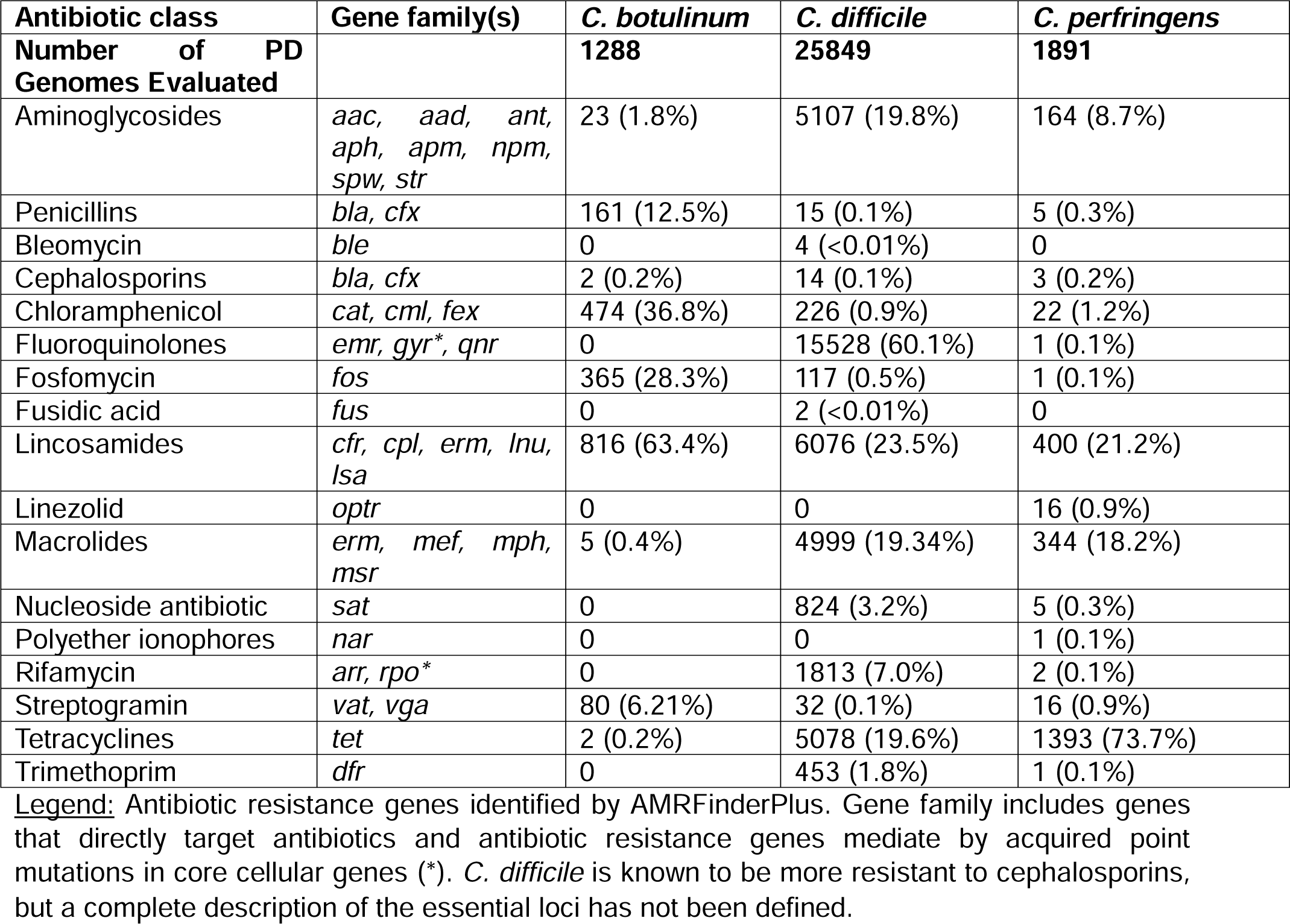
AMRFinderPlus-identified resistance genes in *C. difficile, C. perfringens,* and *C. botulinum*.

In *C. difficile*, fluoroquinolone resistance is the most predicted phenotype with 60.1% (n=15,528) of isolates carrying a corresponding resistance gene or variant allele, particularly in the *gyrA* or *gyrB* DNA gyrases. In contrast, fluroquinolone resistance has not been predicted in *C. botulinum* genomes communicated to date, and in only one strain of *C. perfringens* (1/1,891). The lower rates of fluoroquinolone resistance in these latter two species may reflect true susceptibility but also lack of sufficient genomic content to predict resistance. Continuing research on drivers of fluroquinolone resistance in *C. perfringens* and *C. botulinum* may support future calls of underlying resistance^66^.

Among other antibiotic classes, resistance to lincosamides, such as clindamycin, occurs frequently in *C. botulinum* (n=816, 63.4% of strains), driven primarily by the genes *cfr* (n=811 strains) and *lsa* genes (n=423 strains). In contrast, lincosamide resistance occurs less frequently in *C. perfringens* (n=400 strains, 21.15%) and in *C. difficile* (n=6,076 strains, 23.51%) though clindamycin is a known trigger of CDI. *catA-*driven chloramphenicol resistance also occurred most frequently in *C. botulinum* (n=474, 36.8%), as compared to *C. difficle* (n=226, 0.87%) or *C. perfringens* (n=22, 1.16%). Fosfomycin resistance driven by a *fos* gene was also more common among *C. botulinum* strains at 28.3% (n=365), and rare in *C. difficile* (n=117, 0.45%) and *C. perfringens* (n=1, 0.05%). Notably, fosfomycin and chloramphenicol are used commonly veterinary settings, in poultry and swine^67^, an association which raises interesting questions about exposures that may enhance the acquisition and transmission of resistance genes among *C. botulinum* and other toxigenic species.

The most frequent form of antibiotic resistance in *C. perfringens* was against tetracyclines (n=1,393 strains, 73.7%) primarily via carriage of *tetA*(P) (n=1,384 strains) and/or *tetB*(P) (n=790 strains). Among strains of *C. difficile* tetracycline resistance occurred at lower rates (n=5,078 strains, 19.6%) driven primarily by carriage *tet*(M) (n=4641), while only two isolates of *C. botulinum* harbored resistance (0.16%). Though both *C. perfringens* and *C. botulinum* are associated with farms, the evolution of their AMR repertoires suggests widespread, fundamental differences in routine antibiotic exposures that may relate to habitat and agricultural practices.

Groups investigating drivers of antibiotic resistance should note that the combination of genomic and phenotypic resistance data provide the most robust information to evaluate functions of genomic loci in promoting resistance, and to exercise caution in interpreting resistance gene calls in the absence of phenotypic confirmation. For example, in *C. difficile* AMRFinderPlus has identified plasmid-borne genes that increase strain phenotypic resistance to metronidazole^68^. In contrast, many *van* gene homologs that are involved in *C. difficile’s* cell wall synthesis, promote phenotypic resistance to vancomycin in *Enterococci* and other species, but do not do so in *C. difficile ^47^.* Similarly, chromosomal carriage of a class D beta-lactamase in *C. difficile^69^* was not associated with phenotypic resistance to multiple penicillin, cephalosporin, or carbapenem agents when tested *in vitro ^47^.* The reasons for these differences may relate to differing functions of these genes in *Clostridia* versus other members of phylum Firmicutes, as well as the conditions that induce their expression in *in vitro* versus *in vivo* or *ex vivo* settings.

### *C. difficile* genomic clusters shared among human, veterinary, and environmental reservoirs

*Clostridioides difficile* is an opportunistic pathogen that causes pseudomembranous colitis in humans and other mammalian hosts, commonly after antibiotic disruptions to the gut microbiota^70,71^. *C*. *difficile* infection (CDI) develops with the secretion of toxins (Table 2), particularly the Pathogenicity Locus (PaLoc) toxins *tcdA,* which produces Toxin A (86.2% of deposited strain genomes), and *tcdB,* Toxin B (92.0% of deposited strain genomes), which cause the severe diarrhea and inflammatory responses associated with pseudomembranous colitis, and in very severe cases, cause toxic megacolon. Many epidemic strains, particularly among ribotype 027 strains, harbor the additional binary or cytolethal distending toxin (*cdtAB;* 28.2% of deposited strain genomes), encoded in the unlinked CdtLoc locus. Among toxins, the *tcdB* ADP-ribosylating toxin is most associated with symptomatic disease through its damage of host mucosal surfaces ^72^.

**Table 2:** *Clostridioides difficile* genomic toxin types in NCBI’s Pathogen Detection.

| Toxin genotype | Number (n=25849) | Percent of total |
| --- | --- | --- |
| *tcdB* | 1279 | 4.9% |
| No toxins identified | 1943 | 7.5% |
| *cdtAB, tcdA, tcdB* | 6923 | 26.8% |
| *tcdA, tcdB* | 15270 | 59.1% |
| Others | 434 | 1.7% |

**Legend:** Named toxin types are those with a prevalence above 4%. Others includes 12 predicted toxin types. Data shown for deposited *C. difficile* genomes as of December 2023.

Approximately 59.1% of *C. difficile* strains in Pathogen Detection carry *tcdA* and *tcdB*, while 26.8% carry *tcdA*, *tcdB*, and *cdtAB*, particularly in epidemic strains associated with sequence types ST1 and ST2^72^. Among isolates, 4.9% encode *tcdB* alone, a toxin type found more commonly in infections caused by ST37 isolates^48^. Notably, 7.5% of deposited isolates encode no toxin, representing commensal and environmental isolates that do not cause pseudomembranous colitis. We also note that communicated genomic datasets may skew towards clinically presenting cases given primary contributions from clinical and veterinary surveillance programs.

While CDI is the most common healthcare-acquired infection (HAI), improved infection control practices, antibiotic stewardship programs, and means to implement surveillance programs for *C. difficile* have been shown to be effective in reducing hospital-based rates for CDI ^73,74^. In contrast, the incidence of community-acquired CDI has been on the rise, doubling in the past decade. The dynamics of community infections tie into patient acquisition from additional reservoirs, including from the environment and from animals^58,75^, thus emphasizing the need to approach CD prevention and surveillance from a One Health perspective across clinical, veterinary, agricultural/food-borne and environmental settings to enhance surveillance practices (Figure 2).

**Figure 2:**
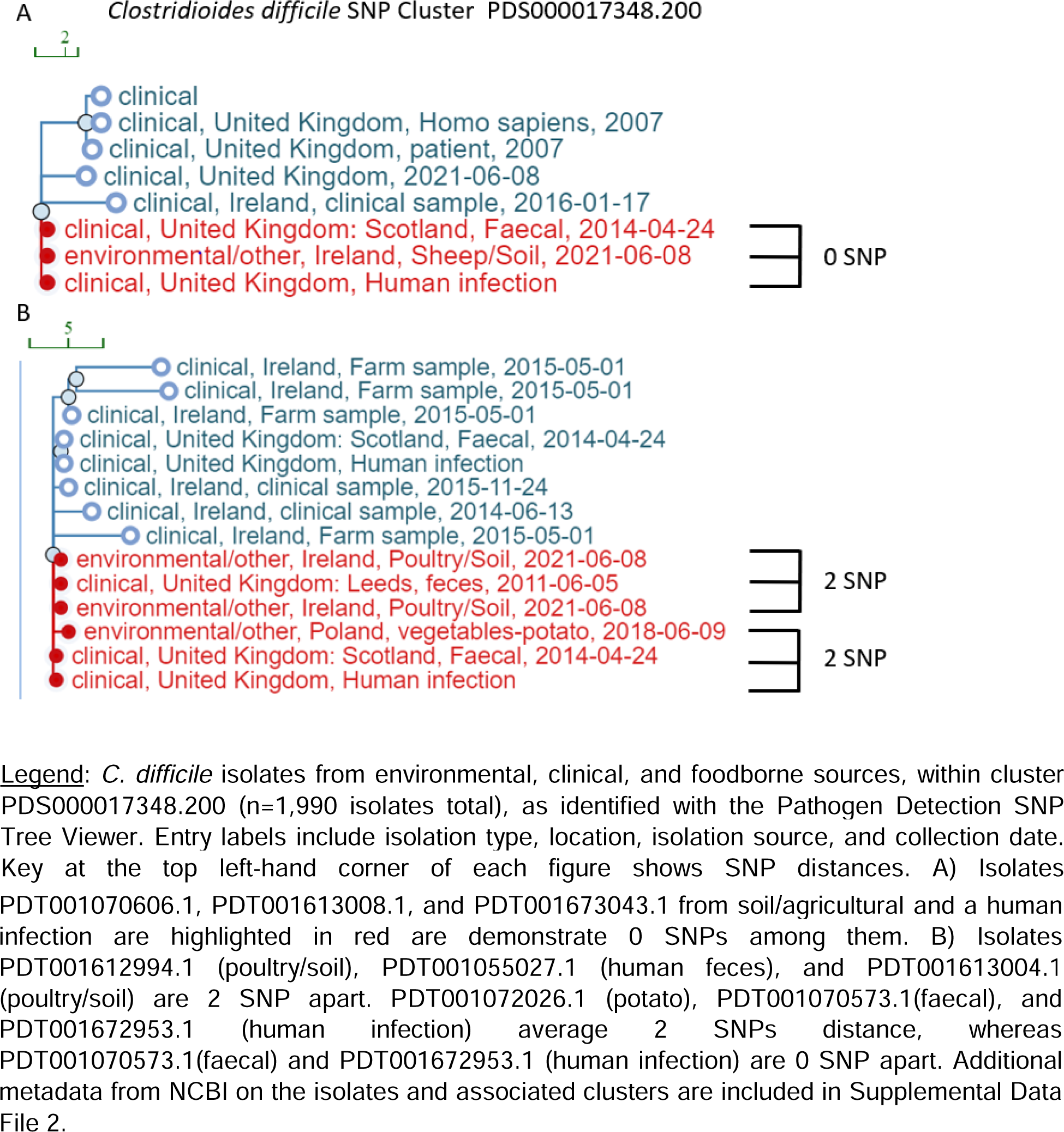
Example toxigenic *C. difficile* genomic clusters from environmental, foodborne, and clinical reservoirs.

The global prevalence of *C*. *difficile* from animals and environmental sources between 2000-2019 ranged widely from <0.5 to 100% detection by source and country ^76^. *C*. *difficile* was identified in farm animals, food sources, natural environments including in compost and soil, and in household settings, including from bathroom toilets and children’s sandboxes. The relationship between and the genomic distribution into 65 sequence types (ST) from human, canine, and environmental isolates in Flagstaff, AZ was also reported for 562 genomes^30^. Eight STs were shared among the three sources with genomic similarities and epidemiologic linkages identified among multiple canine and soil isolates, suggesting direct mechanisms of acquisition and transmission among these reservoirs^76^. In agreement with prevalence observed clinically, ST2 and ST42 were the predominantly observed in canine samples which is likely due to the close interactions shared between the two groups.

### *C. perfringens* genomic clusters shared among foodborne, environmental, and clinical cases

*C*. *perfringens* is the second leading cause of foodborne illness in the US with an estimated 1 million illnesses per year (CDC)^23^. As with other *Clostridial* species, *C*. *perfringens* is ubiquitous and occur commonly in wastewater, soil, and in foodstuffs, with undercooked beef and poultry as the most common causes of foodborne illness. From 1998-2010, 289 outbreaks and over 15,000 illnesses of *C*. *perfringens* were reported to the CDC with 66 outbreaks attributed to beef^77^.

Toxigenic *C. perfringens* (Table 3) carry highly variable and diverse toxin repertoires^78^. Alpha toxin is the most prevalent toxin gene in *C. perfringens*, occurring in 98.6% of deposited genomes (*cpa*), followed by perfringolysin O (*pfoA*; 71.4%), beta2 toxin (*cpb2*; 49.1%), and enterotoxin (*cpe*; 20.2%). Additional toxin genes occurring at <5% prevalence among deposited strains include: *becAB, cpb, cpd, etx, iap/ibp, netB, netF,* and *tpeL*. *C. perfringens* types A and C are most associated with disease outbreaks, with type A associated with foodborne illnesses.

**Table 3:** *Clostridium perfringens* genomic toxin types in NCBI’s Pathogen Detection, as of December 2023.

| Toxin genotype | Number (n=1891) | Percent of total |
| --- | --- | --- |
| *cpa, cpb2, cpe, pfoA* | 101 | 5.3% |
| *cpa, cpb2* | 122 | 6.5% |
| *cpa, cpe* | 139 | 7.4% |
| *cpa* | 234 | 12.4% |
| *cpa, pfoA* | 404 | 21.4% |
| *cpa, cpb2, pfoA* | 547 | 28.9% |
| Others | 344 | 18.2% |

Legend: Named toxin types are those with a prevalence above 5%. Others include 26 predicted toxin types. Data shown for deposited *C. perfringens* genomes as of December 2023.

Genomic analyses of case clusters identified in New York in 2020 leveraged the GenomeTrakr’s GalaxyTrakr pipeline to definitively call transmission chains from contaminated potatoes and gravy with human cases of food poisoning^23^ (Figure 3). Equally important, genomic analyses also successfully ruled out specific isolates as being involved in the case clusters, a factor that can streamline epidemiologic investigations. As more sites contribute genomes over time, the added depth will inform occurrence of sequence types and toxinotypes among areas of surveillance, as well as the extent to which virulence gene exchange may contribute to *C. perfringens* evolution in different settings.

**Figure 3:**
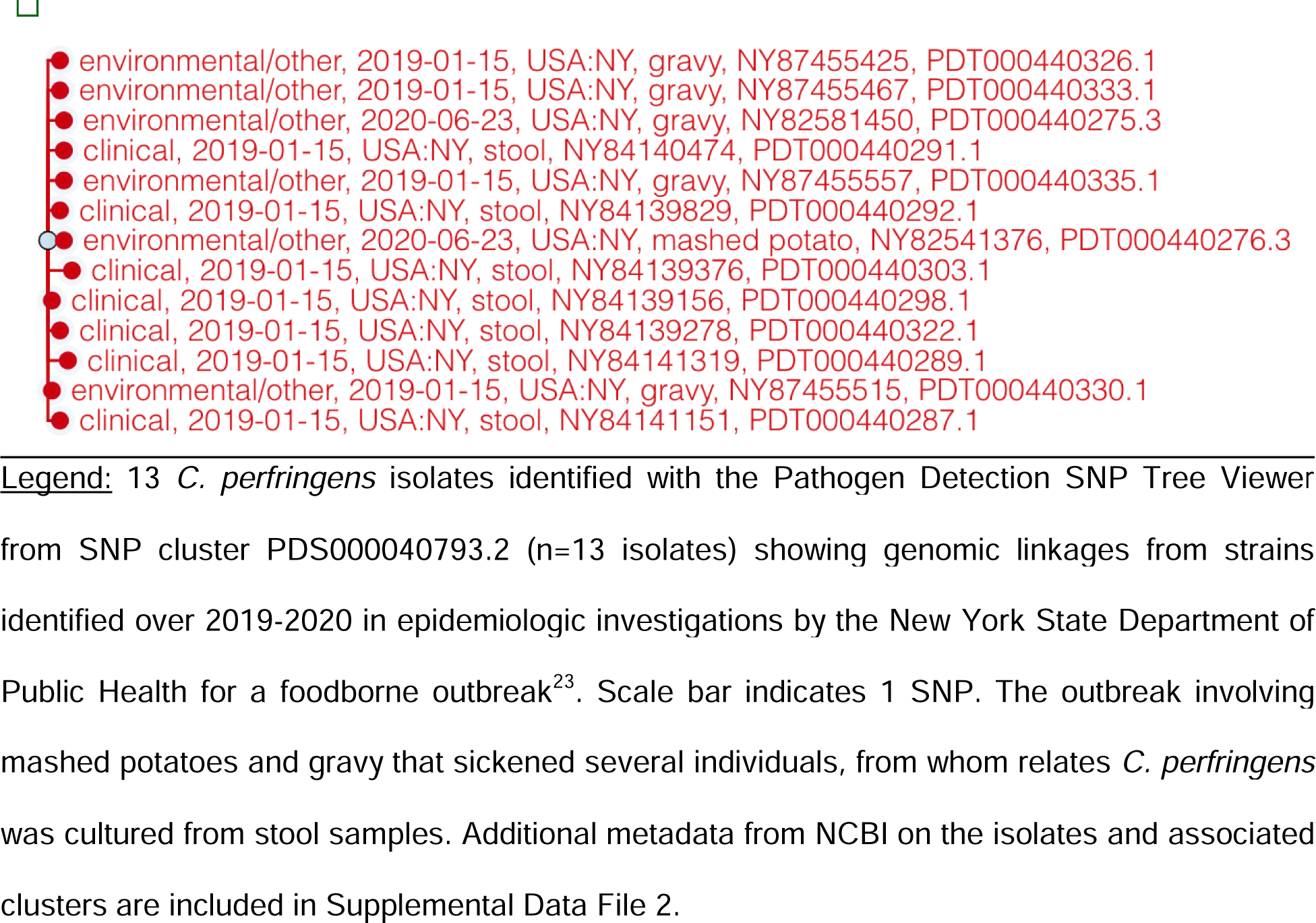
Genome clusters of *C. perfringens* identify foodborne causes of food poisoning.

**Figure 4:**
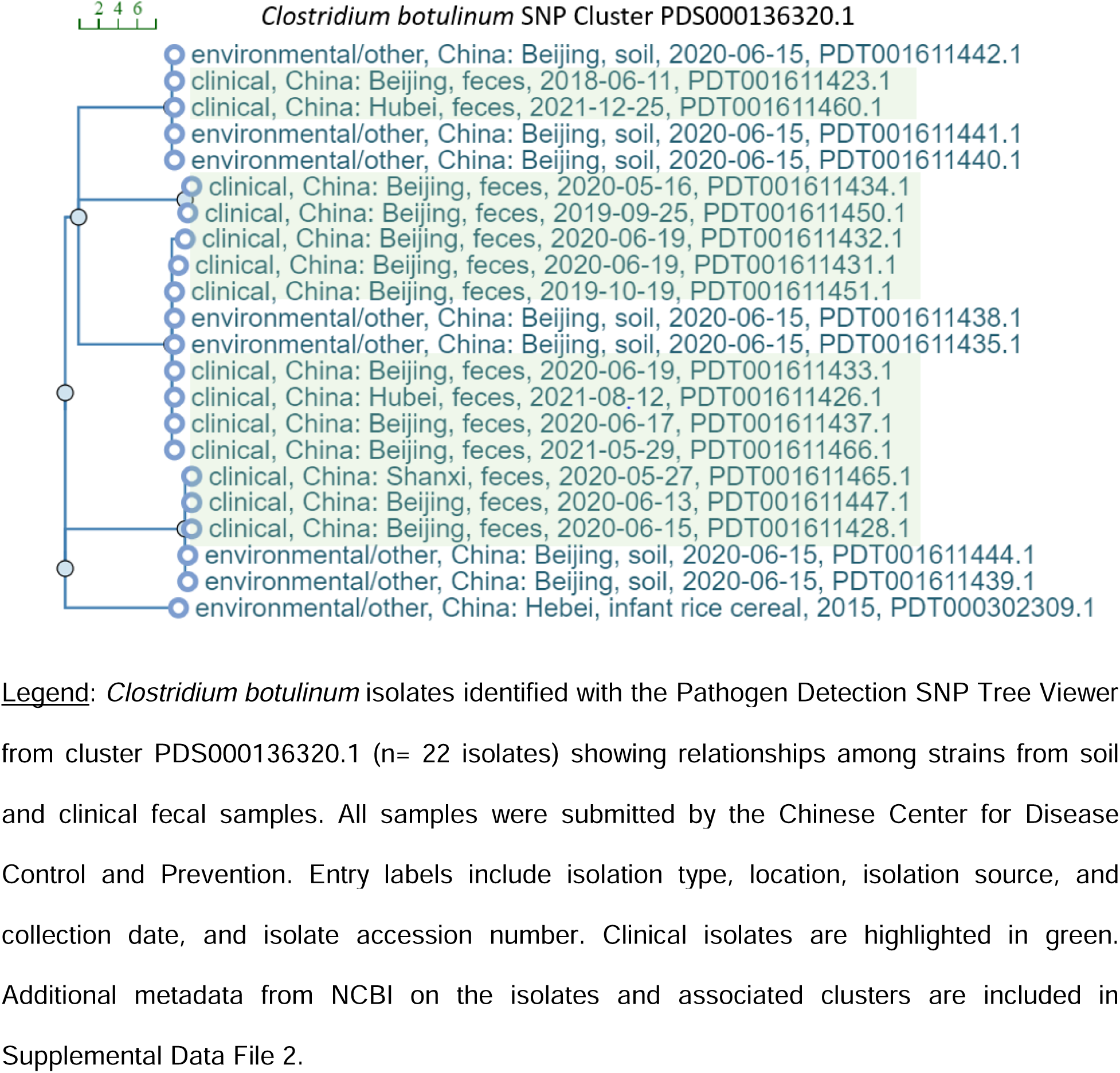
Genomic clusters of *C. botulinum* identify time-linked clusters between foodborne outbreaks and patient cases.

### *C. botulinum* genomic clusters shared among foodborne, environmental, and clinical cases

*Clostridium botulinum* is a mesophilic spore-former that produces botulinum neurotoxins (BoNT) that infect both animals and humans causing severe muscle paralysis^79^. *C*. *botulinum* is categorized into seven toxinotypes, A-G and divided into four main groups, I-IV with 40 recognized neurotoxin subtypes identified through sequencing^80^, information that may inform future PD resources for toxin calling in this species.

Botulism is one of the most lethal foodborne pathogens. Given its severity, surveillance from a One Health perspective is warranted to detect its presence in foodstuffs and zoonotic reservoirs^10^. A total of 10 human botulism and over 50 outbreaks in animals (wild and domesticated) per year were reported in France between 2008-2019 as part of a case-based passive surveillance system^81^. The National Reference Center for Anaerobic Bacteria and Botulism (NRC) monitors human botulism and provides biological confirmation in France. In September 2023, an outbreak of 15 type B botulism cases identified by the NRC in patients from seven countries originated from canned sardines served at a restaurant in Bordeaux, France during the Rugby World Cup^82^. Additional case clusters show linkages between foodborne and patient isolates. The National Botulism Surveillance Summary of 2019 from the CDC reported a total of 215 cases in the US with the majority from infant botulism cases (71%), wounds (19%), and foodborne (10%) ^83^. Toxinotypes A (43%) and B (54%) were the most frequently reported followed by bivalent strains that produce more than one neurotoxin, Bf, Af, and Ba. Type B toxin was reported in nearly 80% of confirmed botulism incidents in Italy between 1986-2015 with most cases associated with the consumption of home-canned products^84^.

A longitudinal cohort of infant botulism type B cases and soil samples were reported in PD by the Chinese Center for Disease and Prevention from Northern China^85^. A total of 22 isolates in cluster PDS000136320.1 (Supplemental Data File 2) demonstrated an average of 25 SNPs with a range of 0-38 SNPs among isolates. Several isolates showed 0 SNPs by PD’s SNP calling methods, including soil isolate PDT001611442.1 and clinical isolate PDT001611423.1 from the Beijing area (Figure 3, Supplemental Data File 2). The studies identified a dominance of ST31 and ST32 *C. botulinum* strains colonizing infected infants^85^, and new associations with soil isolates identified from regional sampling.

### Implementing genomically surveillance programs for toxigenic *Clostridia*

Surveillance programs that include genome sequencing of *Clostridial* isolates with robust epidemiologic analyses provide critical information to aid in the rule-in or rule-out potential outbreak clusters. However, these species present unique challenges given the additional equipment and expertise needed to cultivate obligate anaerobes, including their selective culture from microbially dense samples such as stool or soil, or from food products that may have low pathogen concentrations^86^. While enrichment methods, including heat or alcohol-treatment of samples to enrich for spores can be employed, further specific lab equipment, personnel training, and infrastructure including BSL-2 and in some instances BSL-3 space for *C*. *botulinum* are required to support effective microbiologic cultivation^87^. For genomic-based surveillance, molecular laboratory equipment including thermal cyclers, next-generation sequencers such as the Illumina MiSeq or newer long read sequencing platforms, and high-throughput computing systems with deployed bioinformatic pipelines and personnel with computational expertise and data management are required.

More recently, metagenomic approaches to detect pathogenic *Clostridia* in foodstuffs and in samples harboring complex microbial communities have been used^86^. Untargeted metagenomic sequencing, as well as with methods using *ex vivo* enrichment methods for biologic amplification of target species, has worked successfully with pathogens such as *Salmonella enterica*^88^ and *Listeria monocytogenes^89,90^,* and is amenable for use with toxigenic *Clostridia* given readily available methods to enrich for spores in materials inoculated into pre-reduced anaerobic growth media which can be handled in the absence of anaerobic cultivating equipment required for agar plates. Culture enrichment and next-generation sequencing of primary materials containing pathogens has demonstrated comparable resolution of SNP-based analyses to those form genome sequencing of pure isolates^89,91^. With appropriate validation of these methods, they may be used effectively to provide strain genome-level information, including the potential for multiple strains within a species that may occur within a sample. Sites leveraging these methods, particularly on patient samples, need to incorporate robust bioinformatic methods to remove human DNA that may be sequenced in untargeted metagenomic approaches^92^.

## Summary

Genomic resources for toxigenic *Clostridia* offer unique opportunities to investigate pathogen reservoirs that occur globally and that have major healthcare and economic impact on human and veterinary infections, food safety, and environmental monitoring. The evolving international resources have provided tools to improve our detection and tracking of these pathogens, implement improved surveillance and prevention measures, and support active research and public health efforts to understand molecular aspects of their virulence and transmission. As noted in Figure 1, the genomic repositories for *C. botulinum, C. difficile,* and *C. perfringens,* illustrate global contributions from multiple groups, as well as unmet needs to further develop geographic and reservoir coverage. Methods reducing the complexity for their detection, and to incorporate sequence-based methods, including metagenomic approaches, can expand the range of contributing sites and further One Health efforts to further understand molecular drivers and dynamics regarding target reservoirs and better approaches to prevent, diagnose, and treat infections.

## Acknowledgements

These studies were funded by R01AI153605, R01AI79807, P30DK034854, the Massachusetts Life Sciences Center, and the Hatch Family Foundation (Bry). The work of Jay Worley was supported by the National Center for Biotechnology Information of the National Library of Medicine (NLM), and the National Institute of Allergy and Infectious Diseases, National Institutes of Health. We thank Drs. Bill Kilmke and Mike Feldgarden at NCBI for helpful input and discussion.

